# A global *Anopheles gambiae* gene co-expression network constructed from hundreds of experimental conditions with missing values

**DOI:** 10.1101/2022.01.03.474847

**Authors:** Junyao Kuang, Nicolas Buchon, Kristin Michel, Caterina Scoglio

## Abstract

Gene co-expression networks can be used to determine gene regulation and attribute gene function to biological processes. Different high throughput technologies, including one and two-channel microarrays and RNA-sequencing, allow evaluating thousands of gene expression data simultaneously, but these methodologies provide results that cannot be directly compared. Thus, it is complex to analyze coexpression relations between genes, especially when there are missing values arising for experimental reasons. Networks are a helpful tool for studying gene co-expression, where nodes represent genes and edges represent co-expression of pairs of genes. In this paper, we propose a method for constructing a gene co-expression network for the *Anopheles gambiae* transcriptome from 257 unique studies obtained with different methodologies and experimental designs. We introduce the sliding threshold approach to select node pairs with high Pearson correlation coefficients. The robustness of the method was verified by comparing edge weight distributions under random removal of conditions. The properties of the constructed network are studied in this paper, including node degree distribution, coreness, and community structure. The network core is largely comprised of genes that encode components of the mitochondrial respiratory chain and the ribosome, while different communities are enriched for genes involved in distinct biological processes. This suggests that the overall network structure is driven to maximize the integration of essential cellular functions, possibly allowing the flexibility to add novel functions.

## 1. Introduction

The African malaria mosquito, *Anopheles gambiae sensu strictu* and its sister species *Anopheles coluzzii*, formerly *An. gambiae* S and M forms [1], continue to be major vectors of human malaria-causing parasites in sub-Saharan Africa (e.g., [2, 3]). Even with the first malaria vaccine now approved by the World Health Organization, malaria prevention continues to rely largely on vector control, mainly through insecticide use [4, 5]. Insecticide resistance threatens the efficacy of these approaches [5], requiring insecticide resistance management [6] and new vector control strategies to be designed and implemented (e.g., [7]). Systems biology approaches can help to identify new molecular targets for novel control strategies and provide a global view of the consequences of their implementation on mosquito biology. Systems biology approaches are considered in malaria host-pathogen interactions [8, 9]. They also have been applied to vector biology to determine the evolutionary constraints of the mosquito immune system [10], a critical factor in the mosquito’s ability to serve as a competent vector for malaria parasites [11]. To facilitate systems biology approaches in mosquitoes and building on previous work by MacCullum and colleagues [12], we report here the construction and analysis of a global gene co-expression network (GCN) for *An. gambiae*.

Network analysis has been used widely in different areas of science, including for the construction of GCNs to predict gene function and regulation. A GCN is composed of genes represented as nodes, and significant co-expression of pairs of genes is indicated by links. Links are determined by measuring the co-expression patterns of the transcript data under different conditions [13, 14]. High throughput technologies, such as microarray and RNA-seq, allow measuring simultaneously the expression levels for thousands of genes. Network construction and analysis of gene expression data then provides a system-level view of gene expression relationships, identifying the connection between each pair of genes.

Gene expression data are commonly organized into a gene expression matrix that consists of rows representing *m* genes and columns representing *n* conditions (or samples). To construct a gene expression network, first, a similarity score is calculated for each gene pair. If the expression matrix is complete, the matrix consists of *m* × *n* data points, and each gene has an expression vector of length *n*. Thus, comparison of the expression of any given gene pair is based on *n* number of paired elements between their expression vectors, with a paired element between gene *a* and *b* defined as the expression value pair *a_i_b_i_*, where i is a specific condition, with *i* = [1,2,…, *n*]. The similarity between gene pairs in GCNs is commonly measured by the Pearson correlation coefficient (PCC) [14–16]. The PCC outperformed other means of similarity measures when constructing large gene co-expression networks [17]. Once the similarity is calculated between all gene pairs, a similarity matrix *m* × *m* is assembled, with *s_ab_* elements representing the similarity between genes *a* and *b*. Second, based on the similarity matrix, an adjacency matrix is built, which defines network links using a threshold *T*, with a link existing if the similarity is greater than *T*.

Several studies used the PCC to construct a GCN by setting a fixed threshold to determine coexpression between genes, which is appropriate for homogeneous expression data sets, where (1) expression values were obtained with the same technology, (2) the expression value distribution is comparable across all conditions, and (3) the number of paired elements is similar for all gene pairs [13, 14, 16]. However, neither of these three prerequisites are likely to be met, when assembling a GCN based on a large number of data sets that were obtained with different technologies and under distinct experimental conditions. A striking example of such a heterogeneous gene expression data set is that of *An. gambiae*, which underlies the global expression map constructed by MacCullum *et al*. [12]. This data set does not fulfill any of the three prerequisites detailed above for the following reasons: (1) The types of expression values obtained from distinct technologies and downstream analyses are different [12]. For example, in two-channel microarrays, expression values are expressed as ratios between experimental and control conditions. In contrast, in single-channel microarrays and RNA-seq, intensisty of read numbers present expression values. (2) As a consequence, the distributions of expression values vary widely between the different technologies. For instance, the expression values obtained with RNA-seq are overdispersed, ranging from zero to tens of thousands [18]. (3) Missing values are ubiquitous in experiments. In some data sets, the expression of only a subset of genes was sampled for specific purposes (e.g., the *An. gambiae* detox chip [19]). The fact that biological data sets do not meet these criteria presents a major challenge to network biology in general.

Prerequisites (1) and (2) can be met by applying normalization methodologies, including median shift ([12]) and z-score normalization. However, the missing value problem poses a separate challenge for the following reason. According to [20], under the null hypothesis that two genes are not correlated, the number of paired elements between their expression vectors significantly affects the PCC density distribution. The theoretical analysis shows that the PCC distribution with 50 paired elements has a much lower variance than the distribution with ten conditions, which means that the PCC for any given gene pair tends to be lower when there are more paired elements in a given data set. In the *An. gambiae* expression matrix, the number of paired elements for each gene pair is not identical. Thus, a fixed threshold to determine links introduces a bias, as co-expression of genes with a smaller number of paired elements is favored. One way to overcome this challenge was proposed by Lee *et al*. [15]. In their two-step protocol, initially individual networks are constructed for each homogeneous sub-data set, by calculating PCC similarity matrices and using a fixed threshold to select links. The final GCN is then aggregated from the individual networks, by confirming a link if it exists multiple times across the individual networks.

In this paper, we propose a novel method to construct a GCN for *An. gambiae* genes based on several hundred conditions from different publications and platforms. Specifically, to avoid favoring links between gene pairs with a low number of paired elements, we propose a sliding threshold-based method to construct a global *An. gambiae* GCN. To construct the network, we first apply z-score normalization to produce equal variances and means across all conditions. Second, we compute the PCCs of all gene pairs and divide the gene pairs into 26 different groups according to their number of paired elements. Third, we select links based on a sliding threshold, such that, for each group, only the gene pairs within the top 0.5th percentile of PCCs will be connected by a link. The resulting network, which we name AgGCN1.0, remains robust with random removal of up to 15% of conditions. The AgGCN1.0 has similar characteristics to small-world and scale-free networks, as it contains hub genes that are coexpressed with many other genes. Analysis of the network reveals that both the architecture of the core sub-network and the network communities are based on gene function, supporting the power of the proposed method for GCN construction.

## 2. Method

This section presents a brief overview of the expression matrix, and describes the steps to construct the AgGCN1.0 based on hundreds of conditions [12, 18, 21–48], including (1) data pre-processing, (2) PCC thresholding, (3) edge weight assignment, and (4) final network selection.

### 2.1 Description of the gene expression data set

The data set used for constructing the *An. gambiae* GCN, *Anopheles-gambiae_EXPR-STATS_VB-2019-02*, is based on the data set used by MacCullum *et al*. [12], which was updated by addition of several new conditions. *Anopheles-gambiae_EXPR-STATS_VB-2019-02* was publicly available previously through Vectorbase (vectorbase.org; [49]), however, has since been removed in VEuPathDB, after the merger between VectorBase and EuPathDB [50]. *Anopheles-gambiae_EXPR-STATS_VB-2019-02* is based on the AgamP4.11 annotation of the *An. gambiae* PEST genome, and includes log2-transformed expression values of 13,080 genes across 291 conditions, collected from 35 data sets [18, 21–39, 39–48, 51–56]. Each publication contributed on average of eight conditions to the data set, ranging from 1 [42] to 52 [28].

The experimental methodologies varied widely among publications, exploring gene expression changes using different experimental platforms that sampled either across the entire or various sub-sections of *An. gambiae* transcriptome, using total RNA collected from various life stages, tissues, and physiological conditions. Expression data obtained with single-channel microarrays represented 62% of the data set [22–24, 26–30, 33–36, 39], of which 84% were obtained with the Affymetrix GeneChip^®^ Plasmodium/Anopheles Genome Array [22, 24, 26–28, 30, 33–36]. Expression data for the remainder of the conditions were assessed with either dual-channel microarrays (23% of conditions, [21, 25, 31, 32, 37, 39–44, 47, 52–56]) or by RNAseq (15% of conditions, [45–48, 51]). With regards to life stages, 84% of all conditions were analyzed using samples derived from adults, 91% of those from female mosquitoes, a bias that is easily explained by the fact that only adult female mosquitoes transmit vector-borne disease pathogens [11]. Similarly, tissue-specific analyses focused most commonly on the adult female midgut (33% of the conditions that sampled individual tissues [21, 22, 25, 27, 31, 32, 46, 54]), as it constitutes a major bottle-neck for vector-borne pathogens after being ingested by the mosquito with a blood meal from an infected individual [57, 58]. Likewise, the conditions sampled a variety of physiologies that are integral to vector biology and its control, including blood feeding (9% of conditions, [22, 45, 48, 52]), parasite infection (14% of conditions, [25, 31, 32, 36, 37, 46, 54], and insecticide resistance (6% of conditions, [18, 40–42, 44, 53, 55, 56].

### 2.2 Expression data set pre-processing and normalization

This initial data set was partially conflated as it contained 21 conditions [21, 23, 39] that were combined from other conditions in the same data set (e.g., a conflated condition presented a ratio of two other conditions in the data set). Therefore, these 21 conditions did not represent new data, and thus were removed from further analysis. In addition, one platform, the dual-channel microarray *LIV A. gambiae DETOX 0.25k* [19], interrogated the expression of only 226 genes or 1.7% of the *An. gambiae* transcriptome. The inclusion of the 13 conditions that used the *LIV A. gambiae DETOX 0.25k* microarray [52–56] led to a large number of missing values in the data set, and therefore were excluded from the analyses. After their removal, the final data set (Table S1, in Supplementary Materials https://github.com/KSUNetSE/AgGCN1.0) consisted of gene expression data across 257 conditions collected from 30 publications [18, 21–48, 51].

However, direct comparison of expression values across the 257 conditions in the data set was not possible, as the data distribution varied among conditions, not only due to the platforms used, but also due variations between experimental designs and procedures (Table S4, in Supplementary Materials https://github.com/KSUNetSE/AgGCN1.0/). Fig.1 shows the means and variances of expression values in each of the 257 conditions. The log2 mean gene expression values ranged from −0.9 to 10.9, with means for dual-channel microarray data usually around 0, single-channel microarray data ranging between 4.2 and 10.9, and RNAseq data ranging from −0.9 to 2.5 (Table S2, in Supplementary Materials https://github.com/KSUNetSE/AgGCN1.0/). The variance of log2-transformed expression values ranged between 0.01 and 16.5, with no correlation between mean and variance across the data set or data obtained by RNAseq. However, expression data obtained with dual-channel microarray platforms showed a strong positive correlation between expression mean and variance across conditions, while data from single-channel microarrays showed a strong negative correlation. This negative correlation can be largely explained by the data characteristics of individual dual-channel microarray platforms. Data obtained with both the OXFORD Anopheles gambiae Agilent 13k v1 and the Agilent A. gambiae 020449 44k v2 microarray had low means and high variance within conditions, data obtained with the Affymetrix GeneChip^®^ Plasmodium/Anopheles Genome Array showed means and variances in the middle range, and those obtained with the ND Anopheles gambiae Nimblegen 65k v1 microarray had high means and low variance. These observed differences in means and variance across experiments would result in gene expression correlation based on experimental design rather than on underlying gene regulation.

**Fig 1:**
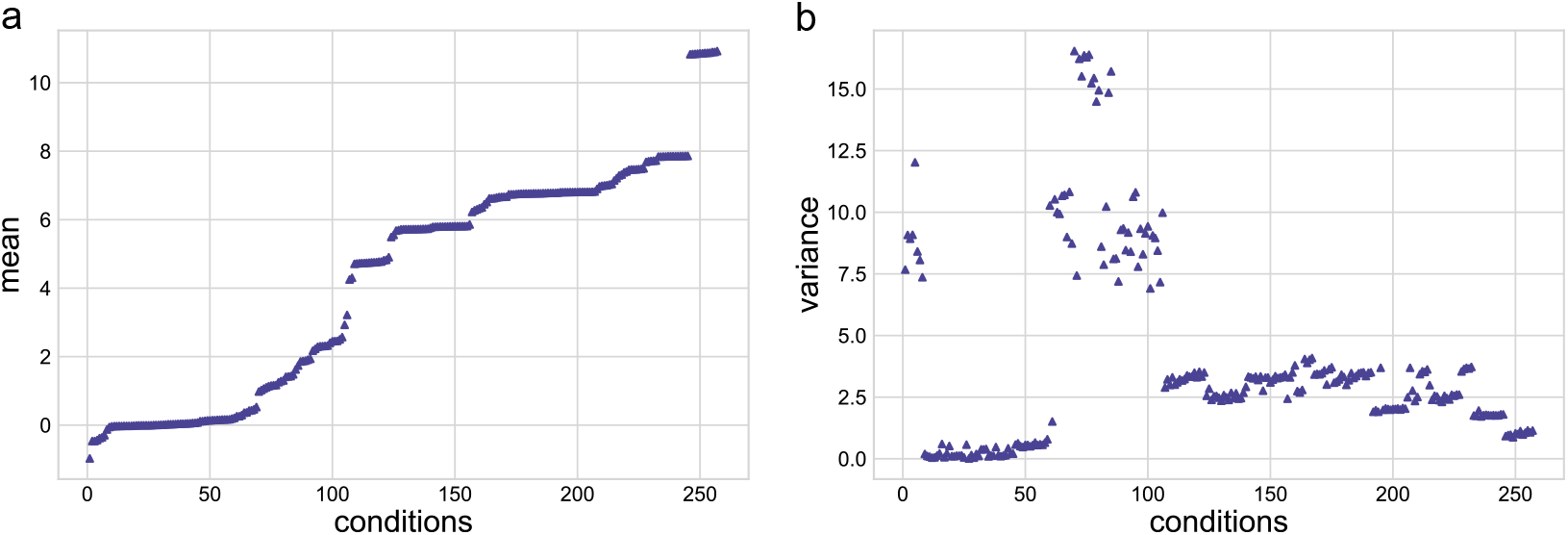
Mean and variance distributions of gene expression values in the data set. Means of expression values for each condition are shown in increasing order (a). The corresponding variance of each mean is shown in (b).

To equalize the means and variance of the data across all conditions, we performed a normalization step using the z-score, which is expressed as follows

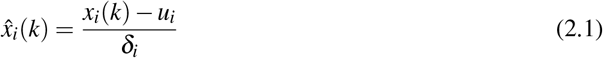

where 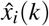 is the normalized expression value of gene *k* under condition *i, x_i_*(*k*) is the raw expression value of gene *k* under condition *i, u_i_* and *δ_i_* are the mean and the standard deviation of the expression value for all values in condition *i*, respectively.

Another property of the *An. gambiae* gene expression matrix was the highly variable length of the expression vectors between all possible gene pairs (Fig. 2). The distribution of paired elements across the entire z-score-normalized data set was highly heterogeneous, displaying a bimodal distribution with a gap between 118-139 paired elements. This gap is explained by the properties of the platforms used to obtain the transcription data. The transcripts of 2,813 annotated genes in the data set were not represented on the Affymetrix GeneChip^®^ Plasmodium/Anopheles Genome Array, which was used to analyze gene expression in 139 out of 258 total conditions [22, 24, 26–28, 30, 33–36]. Therefore, each of these 2,813 transcripts in the data set had an expression vector length that is ⩽118, and the 10,433 transcripts represented on the array had a vector length ⩾139. While z-score normalization addressed the differences in means and variance of the expression matrix, the variable paired element length remained and had important implications on edge selection and edge weight assignment, which are discussed below.

**Fig 2:**
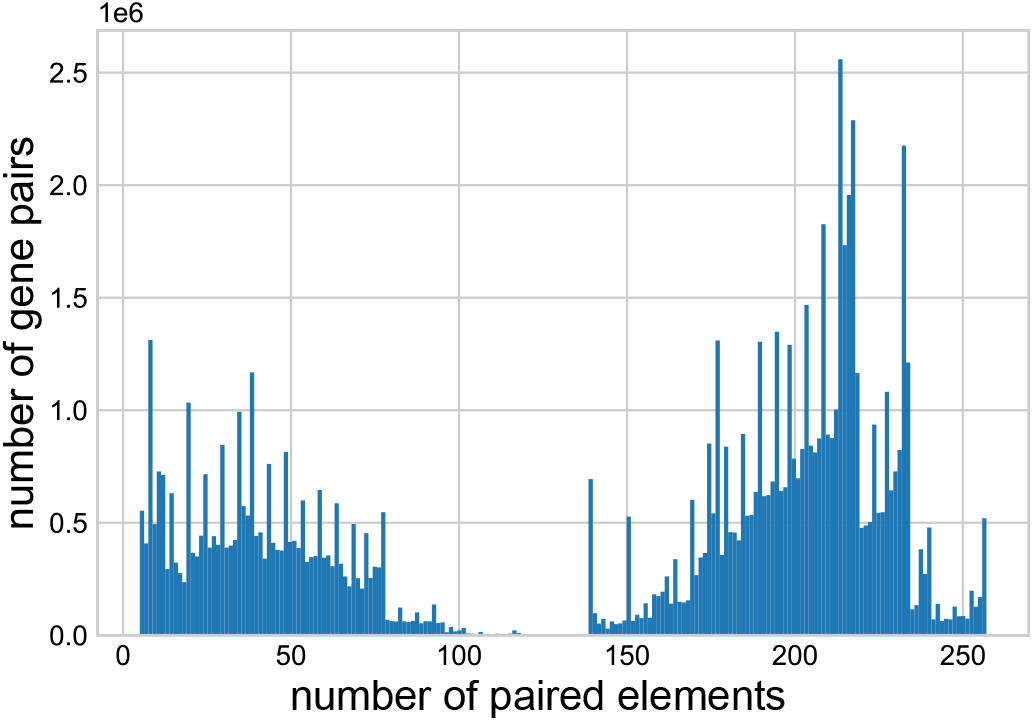
The distribution of the number of paired elements of all the gene pairs.

### 2.3 Edge selection based on a sliding Pearson correlation coefficient threshold

To determine the edges of the AgGCN1.0, we used the Pearson correlation coefficient (PCC), which is a co-expression measure used commonly to detect edges for nodes in gene co-expression networks [15, 20, 59]. We calculated PCCs for all possible gene pairs in the data set, and constructed an initial gene expression correlation matrix, which included all PCCs. From this matrix, we removed the PCCs from all gene pairs whose expression vectors had a paired element length of ⩽4.

The PCC *r_xy_* was calculated as

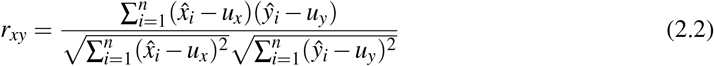

where *n* is the number of paired elements (conditions) between gene *x* and *y*, 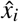 is the normalized expression value of gene *x* under condition *i, u_x_* is the mean of gene *x* across all paired conditions, 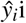 is the normalized expression value of gene *y* under condition *i, u_y_* is the mean of gene *y* across all paired conditions.

From Eq. (2.2), it is apparent that the range of the PCC is [−1, 1]. It is assumed that the two vectors are negatively and positively correlated when *r_xy_* close to −1 or 1, respectively. If the two node-vectors are assumed independent, the density function of PCC is then expressed as

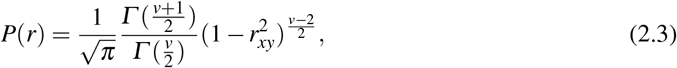

where *Γ* () is the gamma function, and *v* = *n*–2 is the degrees of freedom.

Fig. 3 shows the probability density distribution of the PCC as a function of paired element length. The standard deviation of PCCs decreases with an increase in the number of paired elements. As an example, selecting the top 3rd percentile of all PCCs at a given paired element number lead to a threshold PCC of ⩾0.61 for ten paired elements, and a threshold PCC of ⩾0.12 for 250 paired elements (Fig. 3).

**Fig 3:**
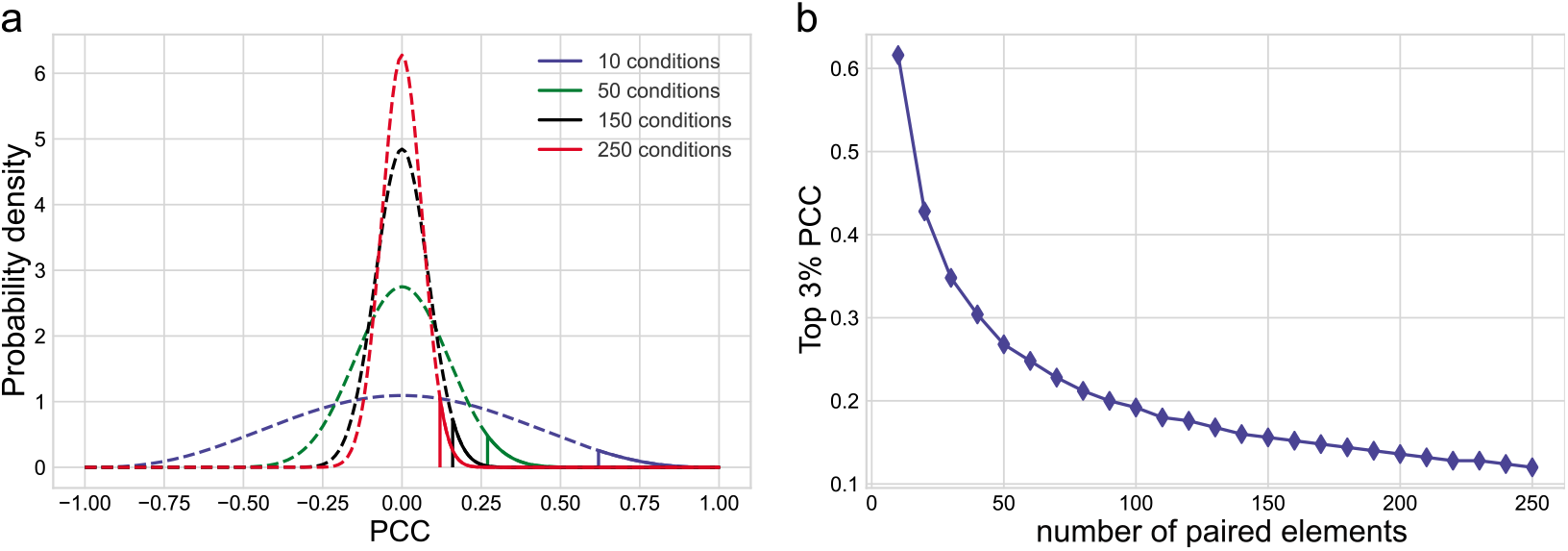
Comparison of probability density function of PCCs with 10, 50, 150 and 250 paired conditions. (a) Probability density functions of PCC, where the vertical lines are the top 3% thresholds. (b) The top 3% PCC with respect to the number of paired elements.

In GCN construction, an edge between a gene pair is included if their expression vectors are deemed to have a *high* PCC, most commonly using a fixed threshold, such as 0.8. However, a fixed threshold is only valid when the number of paired elements in the expression matrix is constant. Given the heterogeneous distribution of paired element length in the *An. gambiae* expression matrix, a fixed threshold of PCC would favor edges between gene pairs with fewer paired elements and not capture the edges with larger numbers of paired elements, which arguably have stronger experimental support.

To avoid this problem, we selected gene pairs among the top percentages of PCC values and used a sliding threshold based on the number of paired elements. To do so, we divided the PCCs into 26 groups according to the interval of the paired element length values, i.e., [4, 10], [11, 20], [21, 30], …, [251, 257]. The scattered points in Fig. 4 show the top 0.5%, 1%, 2%, and 3% of PCCs of all gene pairs in the 26 intervals. To assign a threshold of correlation between the expression vectors of any given paired element length, we used a curve that fitted the scattered points. The gene pairs with PCCs above the fitted curve were assigned edges in the GCN. The equation for the curve is

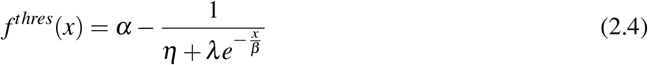

where *α, η, λ*, and *β* are the four parameters that were fitted, and *x* is the number of paired elements. This equation provided a good trade-off between the accuracy of the fitting and the number of parameters to estimate. We optimized the four parameters *α, η, λ*, and *β* iteratively, by (1) fixing parameters *λ* and *β*, and optimizing *α* and *η*, and (2) fixing *α* and *η*, and optimizing *λ* and *β*. We evaluated the performance of the four parameters by maximizing the coefficient of determination, R^2^. The solid lines in Fig. 5 show the fitted curves for the top 0.5%, 1%, 2%, and 3% points, and the optimized parameters *α, η, λ*, and *β* are shown in Table 1.

**Fig 4:**
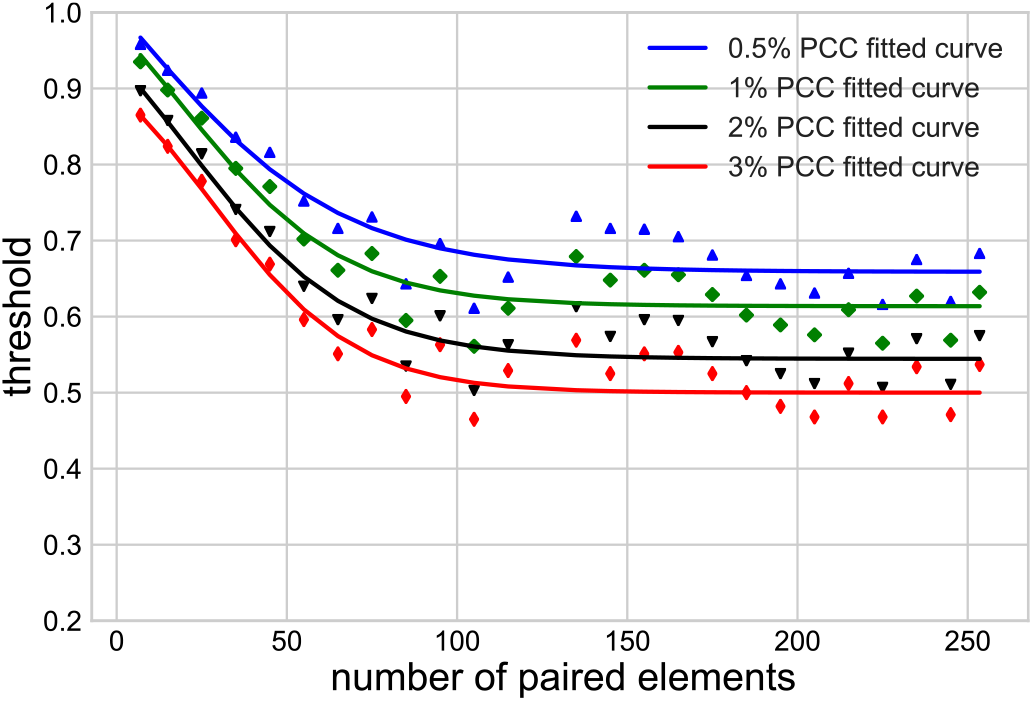
Top 0.5%, 1%, 2%, and 3% PCC points with respect to different numbers of paired elements. Scattered points are the top percentages computed from the data set, solid blue lines are the fitted curves.

**Fig 5:**
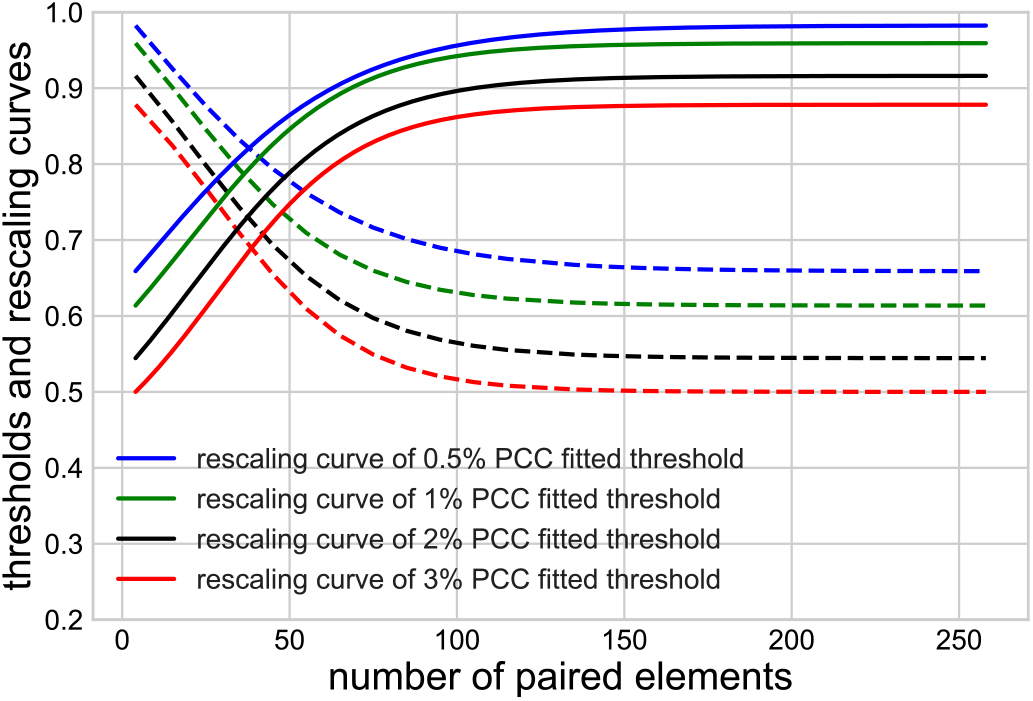
Edge weight rescaling. Sliding PCC-threshold curve (dashed lines) and rescaling curves (solid lines).

**Table 1:** Optimized parameters

| Parameters | $\alpha$ | $\eta$ | $\lambda$ | $\beta$ |
| --- | --- | --- | --- | --- |
| 0.5% | 1.28 | 1.61 | 2.0 | 30 |
| 1% | 1.14 | 1.90 | 4.3 | 23.7 |
| 2% | 1.10 | 1.80 | 4.3 | 24.0 |
| 3% | 1.00 | 2.00 | 7.5 | 21.2 |

Edges were included in the network if their PCC (1) was above the sliding threshold, and (2) had a p-value smaller than *δ/m*, where *δ* and *m* are the significance level equal to 0.05, and the number of gene pairs of each interval, respectively (Bonferroni-correction, [60, 61]).

### 2.4 Edge weight assignment

With the goal to provide a more informative network structure that identifies the relative strength of co-expression between gene pairs, we constructed a weighted network. To do so, we assigned a weight to every detected edge, representing the similarity level of co-expression based on PCC and the number of paired elements. As the edges detected with fewer paired elements tended to have higher PCCs than the edges with more paired elements (Fig. 4), we rescaled the PCCs to equalize the means and medians of the PCCs across the 26 intervals. Fig. 5 shows the PCC-rescaling curves for the sliding thresholds. The dashed lines are the PCC-sliding threshold curves represented by Eq. 2.4, and the solid lines are the corresponding rescaling curves, given by the following equation

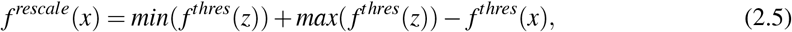

where *z* ∈ [4,257], *z* is the range of the number of paired elements. This curve reduced the weight of those edges that were detected with a smaller number of paired elements. The edge weight was computed as

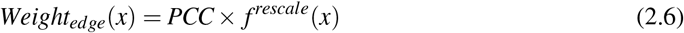

We assigned weights to all edges that were selected based on the sliding thresholds of top 0.5%, 1%, 2%, and 3% PCCs. Fig. 6 confirms that the rescaling curve normalized the means and medians of edge weights across all intervals. As expected, the distribution of the PCC means and medians of the selected edges across the 26 intervals of paired element lengths followed a similar pattern to those observed for the PCC threshold curves (Fig. 6a and Fig. 4). After the rescaling of edge weights using Eq. (2.6), the means and medians of the edge weights in each interval were indeed similar, and thus comparable across the intervals (Fig. 6b), confirming the validity of the rescaling approach.

**Fig 6:**
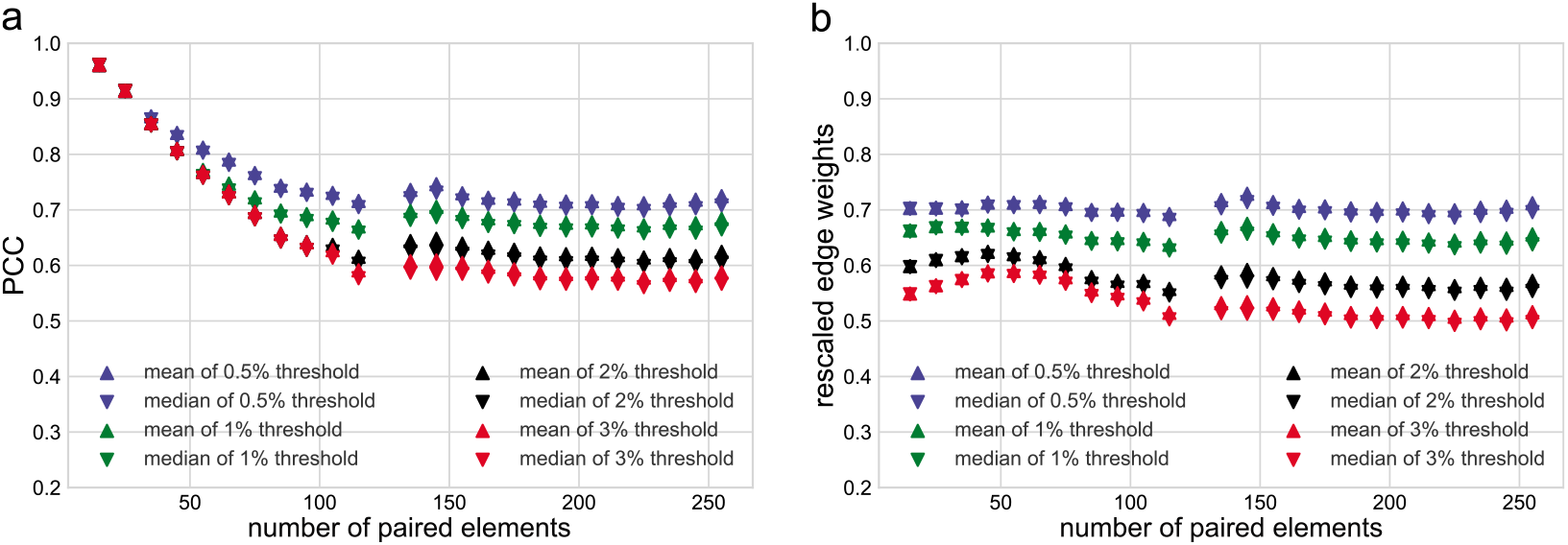
Means and medians of the PCCs above the threshold in each interval. (a) unnormalized means and medians of PCCs. (b) means and medians of rescaled edge weights.

### 2.5 Network selection

To determine the PCC threshold for final network construction, we built and analyzed four networks with the top 0.5%, 1%, 2%, and 3% sliding thresholds as introduced in Section 2.3. Table 2 shows the statistical properties of the four networks. The curated data set contained expression data for 13,080 genes (nodes) annotated in the *An. gambiae* genome. Table 2 shows the number of nodes connected with at least one edge, and the node percentage represents the corresponding percentage of genes present in the network. The LCC is the largest connected component of the network. The network density is defined as

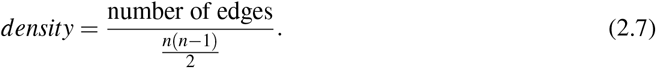

where *n* is the number of nodes in the network.

**Table 2:** Properties of the entire network with different sliding thresholds

| Sliding threshold | 0.5% | 1% | 2% | 3% |
| --- | --- | --- | --- | --- |
| No. of nodes | 11825 | 12290 | 12415 | 12431 |
| Node percentage | 90.4% | 94.0% | 94.9% | 95% |
| No. of edges | 382168 | 685759 | 1307104 | 1896825 |
| Network density | 0.55% | 0.91% | 1.70% | 2.46% |
| No. of connected components | 13 | 4 | 5 | 5 |
| No. of nodes in LCC | 11794 | 12278 | 12401 | 12417 |
| LCC node percentage | 90.2% | 93.9% | 94.8% | 94.9% |
| No. of edges in LCC | 382144 | 685745 | 1307089 | 1896810 |
| LCC density | 0.55% | 0.91% | 1.70% | 2.46% |

All four networks contained more than 90% of nodes, with small node percentage increase with increasing thresholds. In contrast, the number of edges and network density nearly doubled with each threshold increase. Based on these results, we selected the network with the most stringent edge selection criterion (top 0.5% sliding threshold), as the AgGCN1.0 network. This network maintains its structure with a large number of nodes connected with the smallest number of edges. The AgGCN1.0 network, therefore, contains nearly all transcripts encoded in the *Ag. gambiae* genome and shows coexpression between pairs of transcripts only when their expression vectors have very high correlation.

## 3. Methodology verification and robustness test

Next, we validated the methodology of network construction by testing for systematic errors in the procedure and studied the robustness of the AgGCN1.0 network by randomly removing different percentages of conditions from the data set.

### 3.1 Methodology verification

With the construction of the AgGCN1.0 completed, we next verified that the network structure was based on the underlying expression matrix rather than based on a systematic error in the method of construction. To this end, we randomized the expression values under the same condition and reconstructed the network with the method introduced in Section 2 using the following procedure:

Step 1: Randomly reshuffle the expression values for each condition.
Step 2: Compute the PCCs for all of the gene pairs.
Step 3: Use the top 0.5% PCCs fitted sliding threshold to select edges.
Step 4: Repeat step 1 to step 3 100 times.

Fig. 7 shows the distribution of the number of edges of the 100 networks. The AgGCN1.0 network contains 382,168 edges, while the number of edges recovered in the reconstructed networks is smaller or equal to ten. Based on this result, we did not detect a systematic error in the proposed network construction method and concluded that the structure of AgGCN1.0 is indeed based on its expression matrix.

**Fig 7:**
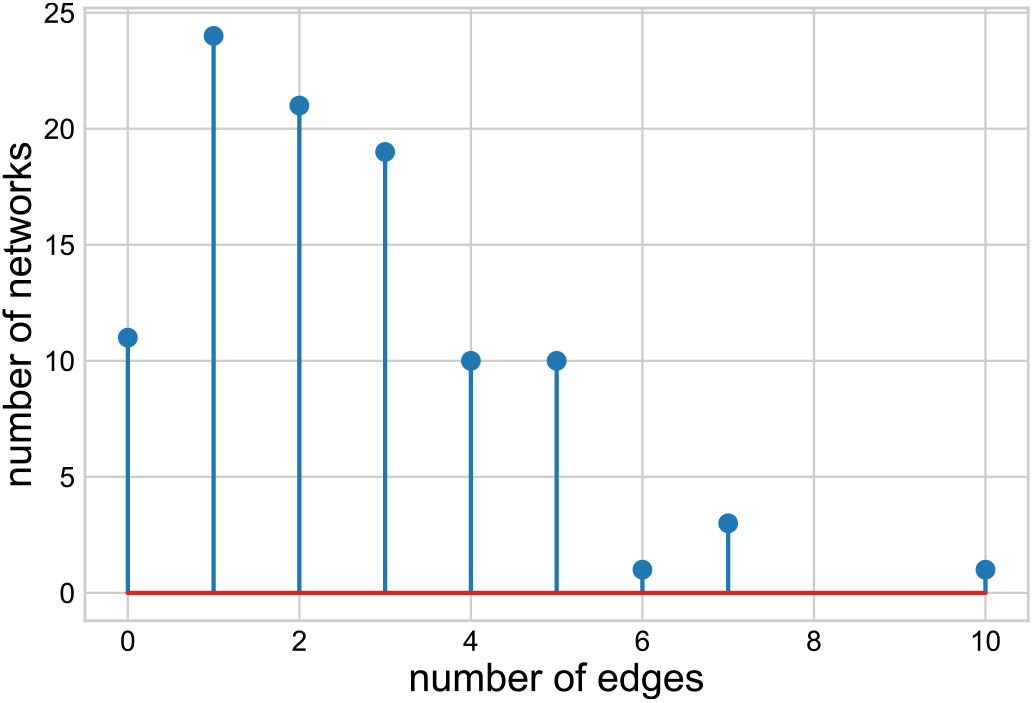
The number of edges from the reshuffled data sets.

**Fig 8:**
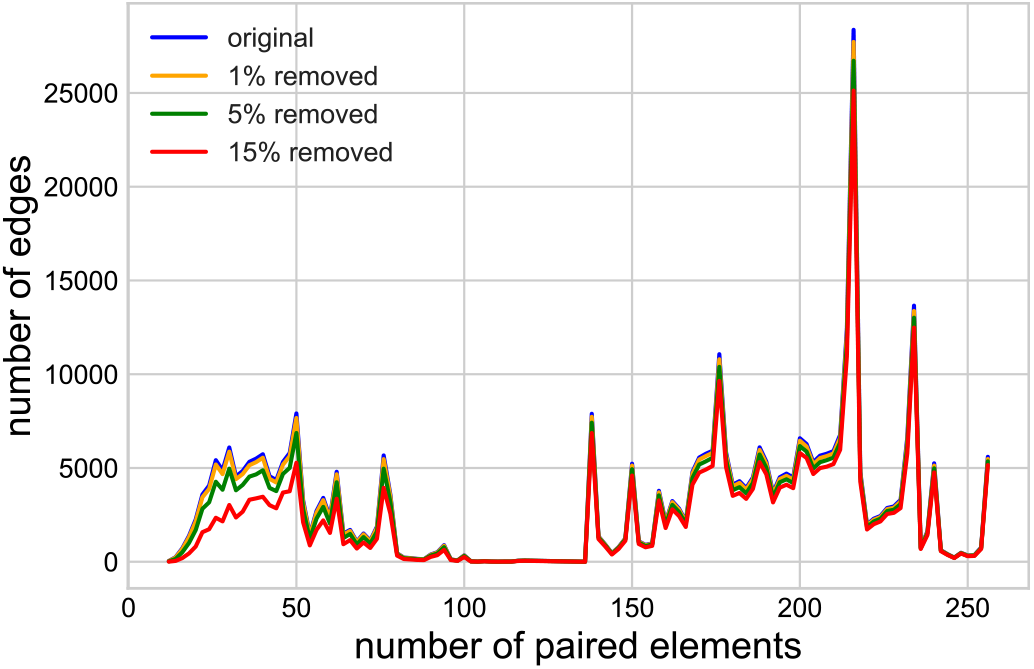
The distribution of remaining edges in the network after removing 1%, 5% and 15% conditions. The horizontal axis is the number of paired elements and the vertical axis is the number of edges.

### 3.2 Robustness test

The AgGCN1.0 network is based on a meta-analysis of expression data obtained from many conditions, and it is unclear how sensitive the overall network structure is to the number of conditions included in the data set. To evaluate network structure sensitivity, we randomly removed an increasing number of conditions and reconstructed the network. The network is considered robust if the edges in the reconstructed networks recapitulate the majority of those in the original network.

Specifically, we randomly and iteratively (100x) removed 3 (1%), 13 (5%), and 39 (15%) conditions and used the top 0.5% PCCs fitted sliding threshold of the AgGCN1.0 to detect edges. Table 3 shows the comparison between the original network and the reconstructed networks. The average number of edges in the reconstructed networks dropped with an increasing number of removed conditions, with a 15% decrease in conditions leading to a reduction of 11.3% in the number of edges. Over 90% of the edges present in the AgGCN1.0 network continued to be detected with 13 conditions removed. However, the number of overlapping edges decreased on average by 22.3% when 39 conditions were removed. To compare more specifically how the removal of conditions influenced edge loss in the AgGCN1.0 network, we determined which AgGCN1.0 edges in each of the 26 paired element length intervals were retained in a sample network that was constructed after removal of (1%), 13 (5%), and 39 (15%) conditions, respectively (8). Overall, edges were largely retained as long as the paired element length was greater than 50. As expected, removal of 39 conditions did not recover edges with fewer paired elements and thus overall low experimental support. Together, these results show that the network construction methodology is robust with respect to random removal of up to 15% conditions. These results also demonstrate that the sensitivity of the AgGCN1.0 structure to the underlying number of conditions is mostly limited to the loss of those edges derived from correlation of expression vectors with few paired elements.

**Table 3:** statistical results of the original and removed networks

| Removed conditions | Gene pairs | Ave. edges | Ave. edges in common (%) |
| --- | --- | --- | --- |
| 0 | 83,781,071 | 382,168 | 382,168 (100%) |
| 3 | 83,768,208 | 381,010 | 366,104 (95.8%) |
| 13 | 83,706,380 | 373,820 | 347,205 (90.9%) |
| 39 | 83,511,993 | 350,306 | 300,939 (78.7%) |

## 4. Analysis of the AgGCN1.0 largest connected component

The AgGCN1.0 network constructed through the method introduced in Sec. 2 using a top 0.5% sliding threshold is composed of 13 connected components (table 2), the largest of which (the LCC) consists of 11,794 nodes and 382,144 edges. In this section, we characterized the LCC network by computing node centralities and detecting its core and communities.

### 4.1 Node centralities

Centralities of network nodes are calculated to detect nodes that can potentially play a critical role in network connectivity, evolution, and dynamics. This general feature extends to GCNs, where node centrality measures have been used successfully to identify genes essential for organism survival [62, 63]. The simplest node centrality is the node degree, which is defined as the number of links incident on a node. When links are weighted, the node degree becomes the node strength, defined as the sum of the weights of all node’s edges. The LCC node strength has an average of 45.7 and spans a wide range of values, included between 0.65 and 965.4. To better understand the characteristics of the AgGCN1.0 LCC, we compared its centralities with the corresponding centralities of two networks generated using the Erdos-Renyi (ER) [64] and the Barabasi-Albert (BA) random network models [65]. By design, the ER and BA networks had the same number of nodes and a similar number of links as the LCC (Table 4). Since the AgGCN1.0 LLC is a weighted network with weights approximately in the interval [0.65, 0.97], in the generation of the two random networks, we assigned weights to the links uniformly at random from the interval [0.65, 0.97].

**Table 4:**
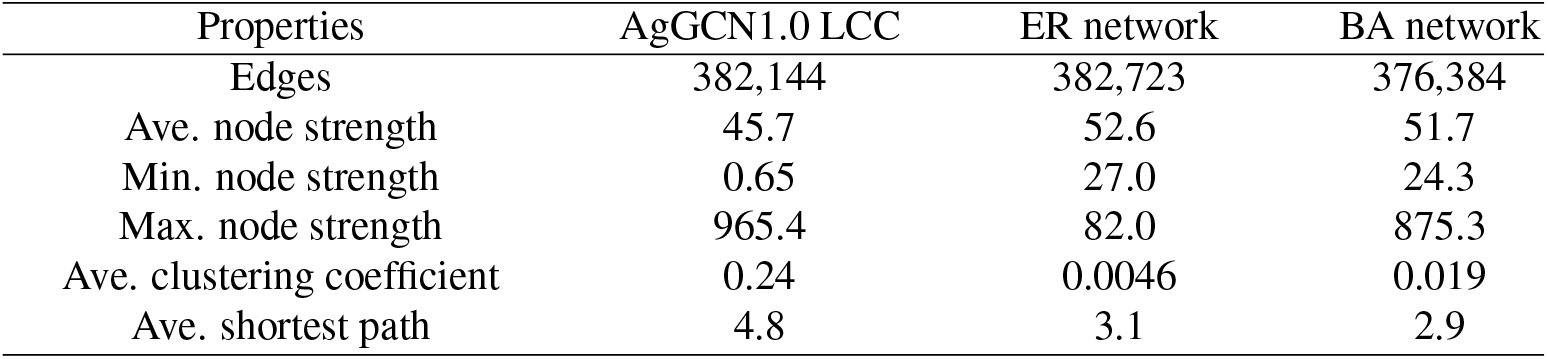
Topological properties of the co-expression network

Table 4 shows the average, minimum, and maximum node strengths of the three networks. While the average value remained similar across the three networks, the range of variability of the ER network was much smaller than those of the two other networks. The similarity of the LCC network with the BA network was also confirmed by comparing the corresponding node strength distributions (Fig.9). These distributions were similar, both showing the presence of *hubs*, i.e., nodes with very large strength values. The presence of hubs is a characteristic of many real networks and provides robustness to the structure with respect to random perturbations of nodes. To measure quantitatively how well the AgGCN1.0 network satisfied the scale-free property, we adopted the model fitting index *R*^2^ on the log-log strength distribution, obtaining values of 0.884 for the AgGCN1.0 LCC and 0.967 for the BA network. The latter value is close to one because the BA network is scale-free by construction. This numerical test confirmed that the AgGCN1.0 LCC can be considered approximately a scale-free network.

**Fig 9:**
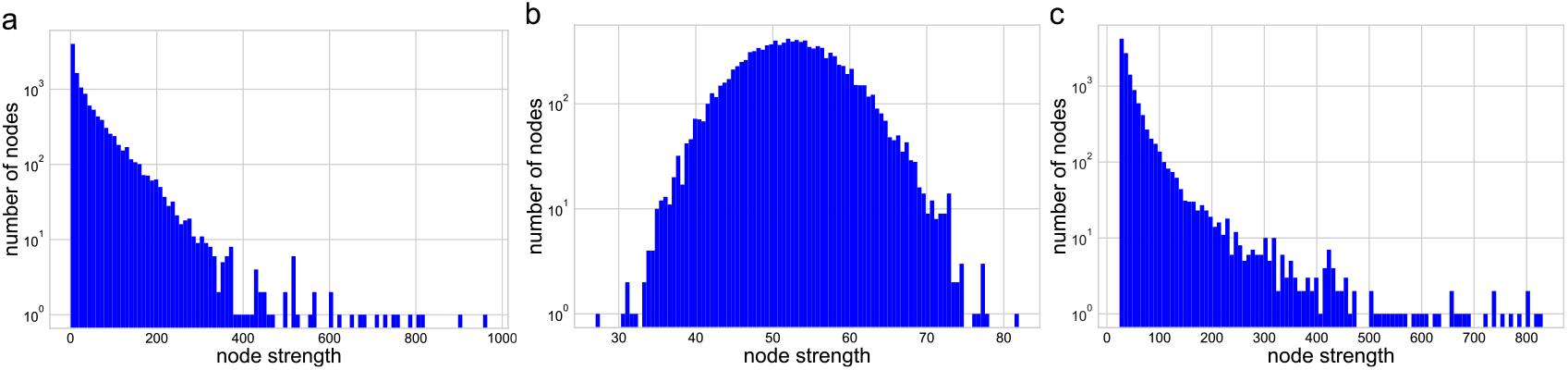
Node strength distribution of the network. (a) The strength distribution of the AgGCN1.0 LCC network. (b) The strength distribution of the ER random network. (c) The strength distribution of the BA random network.

Two other properties of the three networks are computed and compared in Table 4, namely the average clustering coefficient and the average shortest path length. While the clustering coefficient shows how well connected are the neighbors of a given node, the average shortest path length is a measure of the distance of node pairs in the network. The AgGCN1.0 LCC had a much higher clustering coefficient than the ER network and the BA network, which is typical of real networks with communities. In computing the classical shortest path, a small edge weight represents a shorter distance between two nodes. In contrast, in the AgGCN1.0 LCC, higher edge weights represent closer connections. Therefore, the reciprocal of the edge (link) weight is used as the link length to calculate all shortest paths. We then used these shortest paths to compute the average shortest path length, betweenness, and closeness. In Table 4, the average shortest path length of the AgGCN1.0 LCC is compared with that of the ER network and the BA network. The average shortest path of the AgGCN1.0 LCC was only slightly longer in comparison. The two random network models are characterized by the small-world property, which means that there is a path between a pair of nodes that involves only a few short edges and the clustering coefficient is not small. Overall, we can conclude that the AgGCN1.0 LCC also presents some small-world characteristics. Many biological networks, including GCNs show some degree of this property (e.g. [66–68], and it may reflect an evolutionary advantage of such a structure. One possibility is that small-world networks are more robust to random perturbations than other networks and this would provide an advantage to biological systems that are subject to damages, such as gene mutations.

Other centrality measures computed in this analysis, in addition to the node strength, are the following three [69]:

- Eigenvector centrality: the centrality of a node is determined by the entry of the eigenvector corresponding to the largest eigenvalue of the adjacency matrix representing the AgGCN1.0 LCC.
- Betweenness centrality: the centrality of a node is determined by the number of shortest paths that pass through the node itself.
- Harmonic closeness centrality: the centrality of a node is based on its distance to all other nodes. Closeness centrality is the sum of shortest path distance reciprocals of a node to all other nodes. It is calculated as

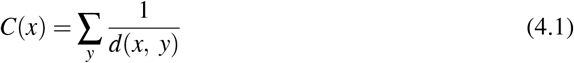

where *C*(*x*) and *d*(*x, y*) are the closeness of node *x* and the shortest path distance between node *y* and *x*, respectively.

Fig.10 shows the four node centralities in the network by visualizing the size of the node proportionally to its centrality. High-strength nodes were distributed in the areas where nodes are tightly connected (Fig.10a, the left and right clusters). However, only nodes in the right cluster also had relatively high node eigenvector centrality, as indicated by their bigger node size (Fig.10b). In contrast, only few nodes had a high betweenness centrality (Fig.10c), while many nodes had a similar harmonic closeness centrality (Fig.10d).

**Fig 10:**
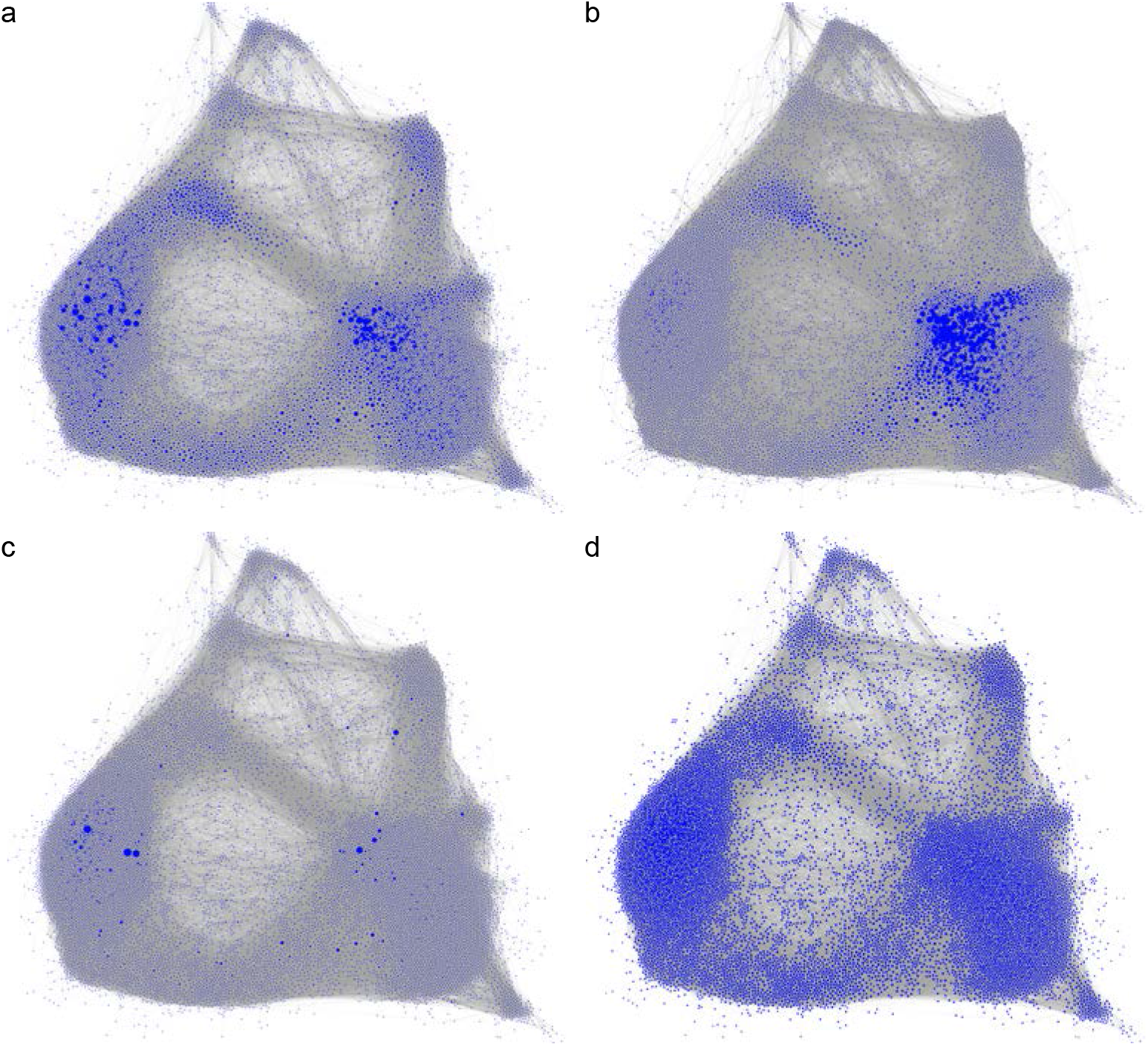
Centrality measures for the AgGCN1.0 LCC. The layout of the network is generated through Gephi with ForceAtlas 2.0 algorithm [70]. (a) node strength centrality. (b) eigenvector centrality. (c) betweenness centrality. (d) harmonic closeness centrality. The size of a node is proportional to its centrality value.

Overall, we did not identify any correlation between gene expression level and either of the four centrality measures reported here (data not shown). Thus, these centrality measures can potentially be used as additional node characteristics. However, since the AgGCN1.0 did not contain a single group of nodes that is characterized by high centrality for all four measures, it will be critical to define the gene property to be studied and determine which measure more closely represents such property to detect the key nodes (genes). Additionally, if an evolutionary process would target the top central nodes in the AgGCN1.0 LCC network, these nodes would be different on the basis of the selected centrality, providing a diversity advantage.

### 4.2 Communities

A community in a network is a subgraph that is highly connected internally and loosely connected with other subgraphs. Community detection for the AgGCN1.0 LCC is essential, since the identified communities can help discover the underlying biological processes that shape the network. In recent years, various community detection algorithms have been proposed. According to [16, 71, 72], the Louvain algorithm and the Infomap perform better than other methods, with the extra advantage of low computational complexity. However, the Infomap method tends to cut leaf nodes into isolated communities, which results in numerous tiny communities. For this reason, in this work, we adopted the Louvain algorithm to detect the communities. The Louvain algorithm is a modularity optimization method that hierarchically identifies how nodes are clustered. Modularity measures the difference between the AgGCN1.0 LCC and a random network in terms of community existence. The modularity *Q* of a network is calculated as follows [71]:

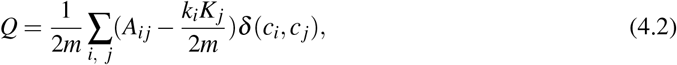

where *Q, k_i_, k_j_, m* and *A_ij_* are the modularity, the degrees of node *i* and *j*, total number of edges in the network and the weight of the edge between node *i* and *j* respectively. The Kronecker delta function δ is equal to 1 if *c_i_* equals *c_j_*, which mean the two nodes are in the same community, while δ is equal to 0 when the two nodes are in different communities.

In the AgGCN1.0 LCC, we detected 15 communities using the Louvain algorithm, as shown in Fig.11. Most of the nodes are included in 13 large communities, while two small communities contain less than 1% of nodes. In the network visualization shown in Fig.11, nodes belonging to the same community were visualized in proximity, due to the algorithm selected to visualize the network. In particular, the layout of the network is based on the ForceAtlas 2.0 algorithm, which produces visual densities that denote structural densities [70]. Visualization of the force-directed layout of the AgGCN1.0 LCC confirmed the existence of well-defined communities. Computing the average shortest path between pairs of communities revealed that communities visualized in proximity are also characterized by a relatively shorter average path length. For example, the average shortest path length between community 11 and community 2 was 3.276, as compared to 4.764 between community 11 and community 15.

**Fig 11:**
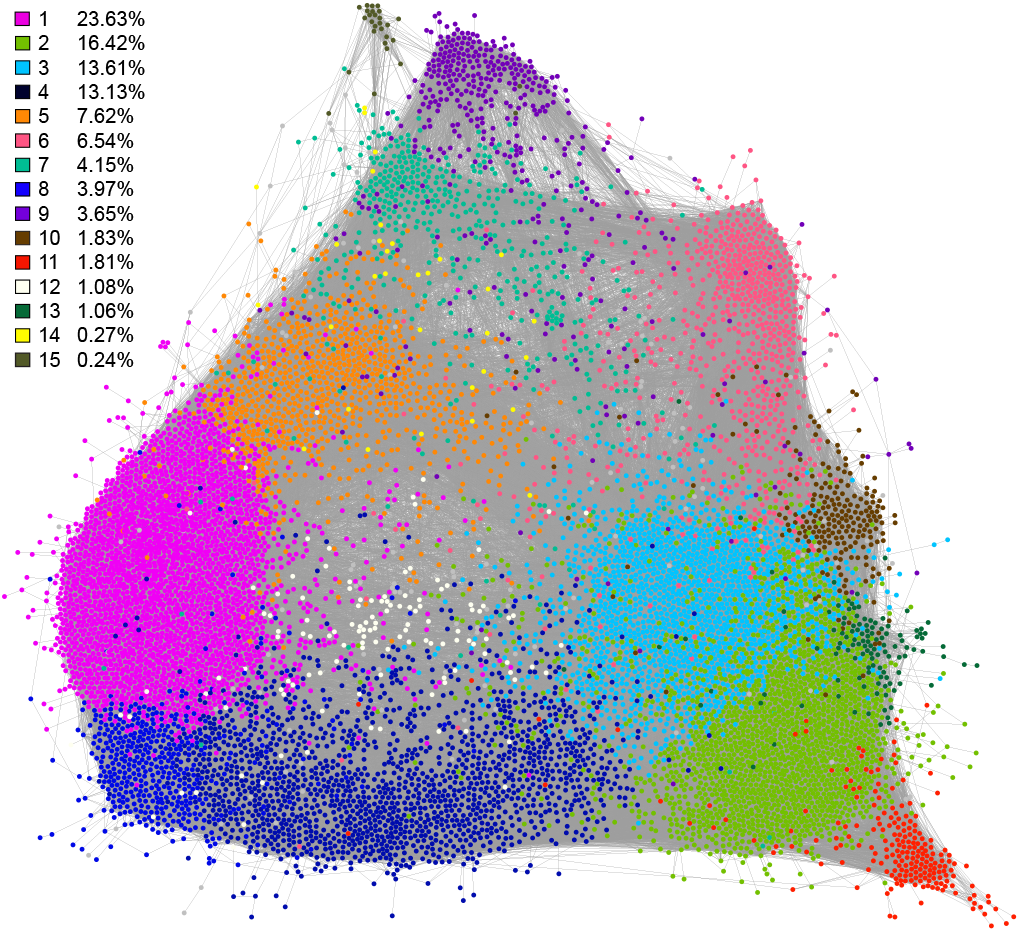
The 15 major communities of the LCC in AgGCN1.0. The network was visualized with Gephi 0.9.2 using the ForceAtlas 2.0 algorithm [70]. Communities are numbered consecutively based on the number of nodes they contain and colored according to the panel on the left.

### 4.3 Cores

The *k*-core of a graph is a maximal subgraph in which each node has at least *k* neighbors after removing nodes with degrees less than *k* repeatedly by starting with *k* = 1 and increasing *k* until no nodes are left in the network. The core of the network is the subgraph obtained with the maximum *k* such that there are still nodes in the subgraph, but with *k* + 1, all nodes are removed. In the case of a weighted network, node strength substitutes node degree, and the definition of coreness needs to be adapted. In the *s*-core decomposition, the *s*-core subgraph consists of all nodes *i* with node strengths *s*(*i*) > *s*, where *s* is a threshold value. We define the threshold value of the *s*_*n*_-core as *s*_*n*-1_=min *s*(*i*), among all nodes *i* belonging to the *s*_*n*-1_-core network. The *s_n_*-core is found by the iterative removal of all nodes with strengths *s*(*i*) ⩾ *s*_*n*-1_. Like k-core analysis, where node degrees are recalculated for every removal, the remaining nodes’ strengths must also be recalculated in the weighted core [73].

The distribution of *s*-core sizes is shown in Fig.12a, while in Fig.12b, the red nodes are the subgraph corresponding to the final *s*-core. If the threshold was above *s*=74.64, all nodes were removed, showing the nodes within the final *s*-core are tightly connected. Green nodes remained at a threshold *s*=48.54, corresponding to the first discontinuous drop distribution of *s*-core sizes (Fig.12a). When *s*=69.20, two separate components were maintained, representing two densely connected parts in the network (blue nodes). After we increased the threshold *s*, the blue node component in the bottom-right disappeared. Note that cores with higher thresholds were included in those with lower thresholds. Nodes in the final core are the ones that remain in the network even when many redundant connections, i.e., many connections to other nodes with equal or smaller strength, were iteratively removed. The final core can be seen as the most critical and internal set of nodes that guarantee network connectivity.

**Fig 12:**
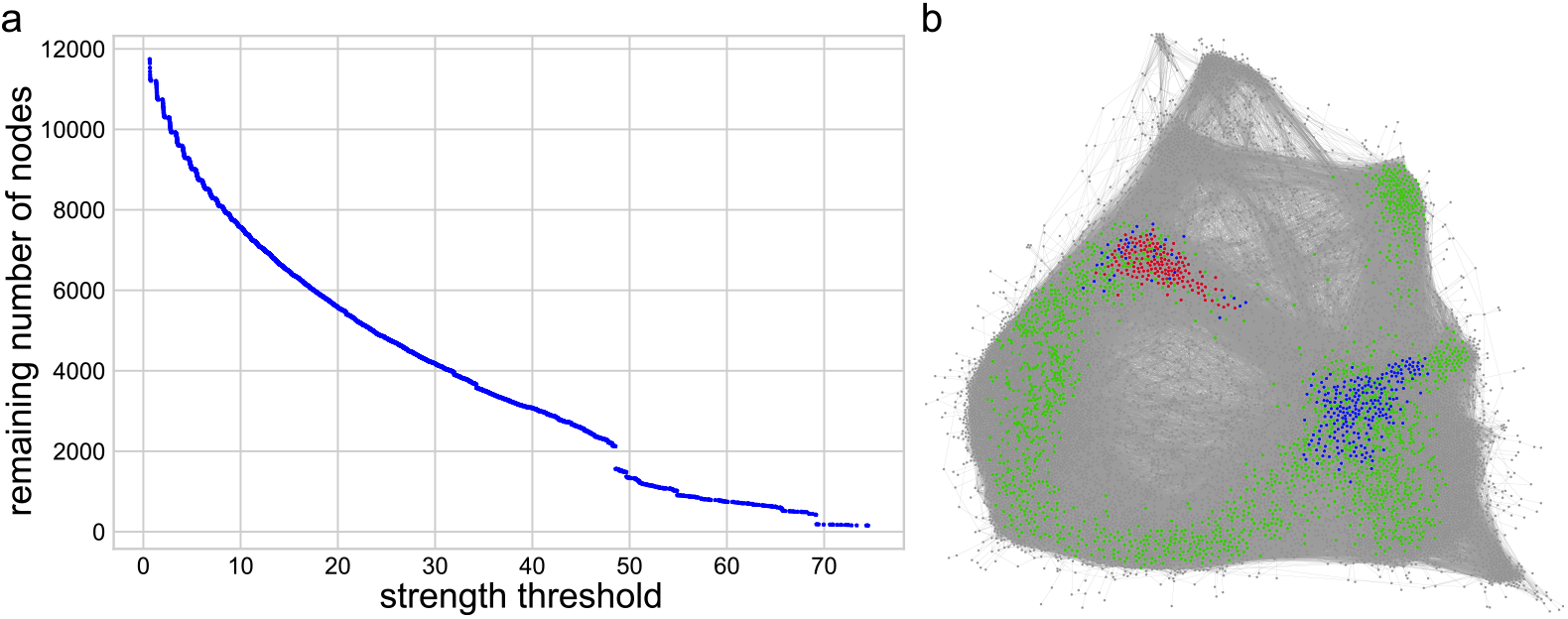
Coreness. (a) The number of nodes left with respect to the *s*-core threshold. (b) Core subgraphs for different strength thresholds. Core *s*=48.54 consists of 2122 nodes (green, blue and red nodes), and core *s*=69.20 consists of 416 nodes (blue and red nodes). The final core *s*=74.64 consists of 153 nodes (red nodes).

The analysis of node characteristics in the final core revealed the following. Core nodes were, on average, highly expressed across most conditions, as compared to all nodes in the LCC, and also sampled across most conditions (⩾230). The centrality measures (strength, eigenvector, closeness, betweenness) of the core nodes were also, on average, significantly higher than those of the LCC. However, the core nodes did not include any of the top central nodes under any of the four centrality measures (strength, eigenvector, closeness, betweenness). Taken together, these results show that the co-expression of core nodes is supported by strong expression across the majority of conditions in the network. In addition, overall gene regulation across the *An. gambiae* transcriptome results in a network structure that at its core maximizes all centrality measures rather than a specific one for each core node. This structure stabilizes the GCN, because a targeted perturbation of the highest centrality nodes would not affect the network core, thus providing another layer of network robustness.

## 5. Network architecture is based on biological function

The analyses of the properties of the largest component of the AgGCN1.0 network identified a smallworld, scale-free network with a small final core and distinct communities. This suggests coordinated behavior of gene expression across a large number of distinct experimental conditions, independent of any single specific condition. Previous studies have identified network architecture of GCNs to be based on gene function, where co-expression modules represented sets of genes that function within the same biological processes (e.g. [74, 75]). To determine whether individual subgraphs of the network were enriched for particular biological processes, we performed a Gene Ontology (GO) and KEGG pathway enrichment analysis in R using topGO [76].

### 5.1 The Network core is enriched in genes required for Oxidative Phosphorylation and Translation

The first subgraph we analyzed for GO and KEGG pathway enrichment was the final core *s* = 74.64, which consists of 153 genes, representing 1.3% of genes that comprise the largest connected component. The top GO categories significantly enriched in the *s*-core group encompass the biological processes of translation and oxidative phosphorylation (Fig. 13, Table S1). Indeed, the *s*-core contains 64 of the 131 genes identified to make up the ribosome of *An. gambiae* (KEGG pathway aga03010), and 38 of the 107 genes that comprise oxidative phosphorylation (KEGG pathway aga00190). We next analyzed GO enrichment in the two larger cores, Core *s* = 48.54 and core *s* = 69.20 (Fig. S2 and 3). Similarly to the final Core 74.64, the GO terms enriched the most belonged to the biological processes of mitochondrial electron transport and translation (Table S1). However, the enrichment was largely due to the presence of genes in the final core, with the larger cores adding less than half of the enriched genes that make up the ribosome and mitochondrial electron transport chain. Given the fundamental need for ATP and proteins for all cellular function, it is perhaps not too surprising that the expression of these genes is most central and integrated across the entire *An. gambiae* transcriptome.

**Fig 13:**
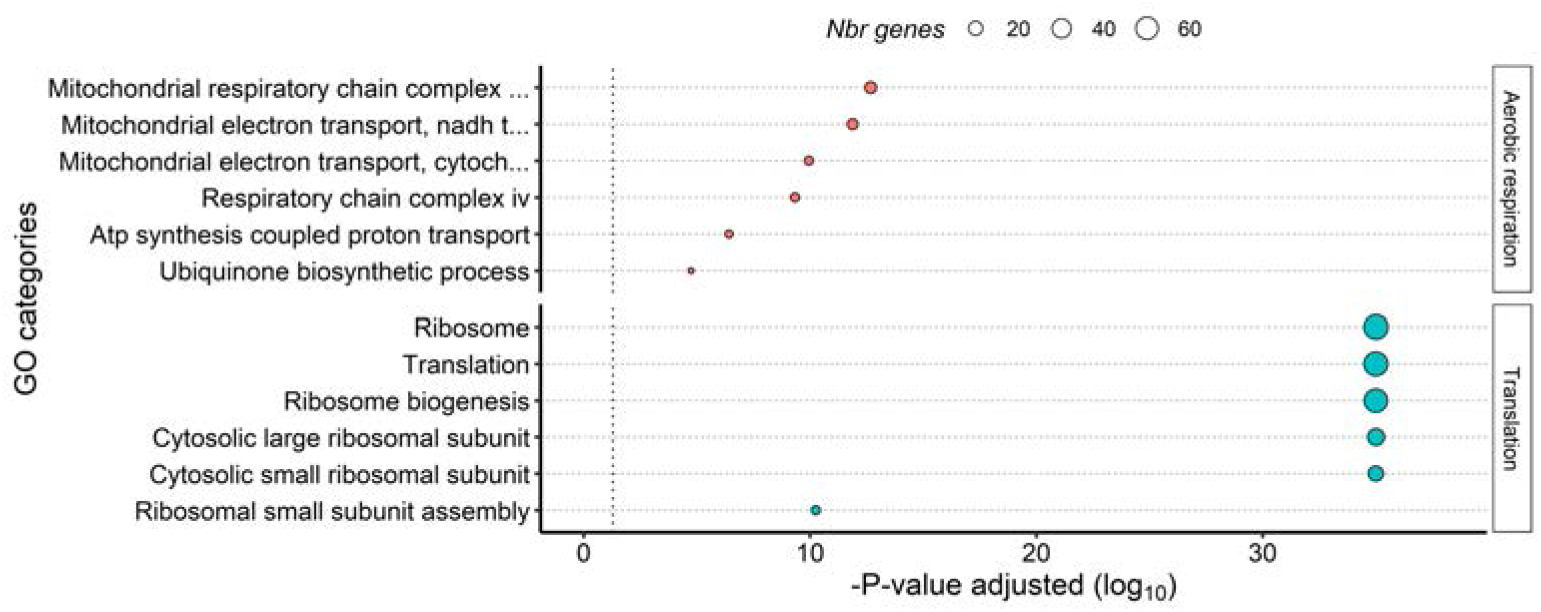
The network core is enriched for genes that function in oxidative phosphorylation and translation. The X-axis represents the statistical significance of the gene ontology (GO) categories (y-axis) after adjustment for multiple testing. Size of the dot is proportional to number of genes in the given GO category.

### 5.2 Network communities are enriched for gene sets with functions in distinct biological processes

We next analyzed the community subgraphs for enrichment of GO terms and KEGG pathways among their annotated genes (results are summarized in Table 5, all data in Table S2, Fig. S3-S17). We found each community enriched for distinct biological processes, which dependent on community, ranged from fundamental cell functions to specialized physiologies executed by specific organs or tissues.

**Table 5:** Nodes distribution in each community and GO term enrichment

| Community<br>(% of network) | Enriched Biological Functions |
| --- | --- |
| 1 (23.6) | DNA repair, Transcription, RNA processing, Translation |
| 2 (16.4) | Neurogenesis/neuronal function, Sensory perception |
| 3 (13.6) | Cuticle metabolism, Cytoskeleton organization, Muscle development |
| 4 (7.62) | Intracellular Signal transduction, Protein phosphorylation, Neuron function |
| 5 (6.54) | Aerobic respiration, Glycolysis/TCA cycle, Redox homeostasis, Translation, Protein processing |
| 6 (4.15) | Neuronal function, Signal transduction, Sensory perception |
| 7 (3.97) | Innate immunity, Lipid metabolism |
| 8 (3.65) | DNA replication/repair, Cell cycle, Cell division, Meiosis/Oogenesis |
| 9 (1.83) | Digestion, Drug metabolism, Chitin metabolism, Transmembrane transport, Innate immunity |
| 10 (1.81) | Chitin metabolism, Lipid metabolism, Proteolysis |
| 11 (1.08) | Cytoskeleton, Axoneme |
| 12 (1.06) | Gluconate shunt, Signaling |
| 13 (1.01) | Cuticle metabolism |
| 14 (0.27) | Proteolysis |
| 15 (0.24) | Salivary gland proteins, Smell* |

\* GO term “smell” due to enrichment of the D7 family salivary gland protein genes, which are odorant binding proteins.

Manual analysis of enriched GO and KEGG terms within each community revealed an additional clustering of individual biological processes that together are required for fundamental cellular functions. For example, Community 1 is enriched for genes with GO terms that broadly fall into the biological processes of DNA repair, Transcription, RNA processing, and Translation, indicating co-expression of genes that belong to the functionally related cellular functions of gene expression (Fig. S3). A second example is Community 5, which contains an overrepresentation of genes with GO term annotations belonging to glycolysis, tricarboxylic acid (TCA) cycle, and Oxidative phosphorylation, and cell redox homeostasis, indicating that that the genes required for oxidative energy production are co-regulated (Fig. S7). In addition, Community 5 is also enriched for genes with function in translation and protein processing, suggesting that the generation of proteins and their turn-over are co-regulated on a transcriptional level (Fig. S7). A third example is Community 8, which is enriched for genes that encode nuclear proteins that function in DNA replication and repair, Cytokinesis, and Cell cycle.

Community 15 presents the most striking example of a set of co-expressed genes that have a highly specialized function, as it is enriched for genes with known function in blood feeding (Fig. S17). About 93% of all genes in Community 15 encode known salivary gland proteins. Indeed, this community contains 51% of all genes identified by Arca et al. as salivary gland protein genes [77]. Furthermore, Community 15 is enriched specifically in salivary gland proteins that are expressed in adult female salivary glands, including the majority of the D7 family, all members of the SG1 family and SG7/SG7-2 family, as well as gVAG (g ambiae Venom AllerGen), apyrase and 5-nucleotidase. In contrast, salivary gland protein genes expressed in male and female salivary glands, including salivary gland amylase, maltase, members of the SG2 protein family, SG9, and the gene encoding the 55.3 kDa salivary gland protein are located in community 9.

Community 7 presents a second example of a set of co-expressed genes that have a highly specialized function, as it is highly enriched for genes encoding proteins with known function in innate immunity (Fig. S. 9). The *An. gambiae* genome encodes 347 canonical immunity genes belonging to the immune modules of recognition, modulation, signal transduction, and effectors [78]. Of these, 23.6% (82 genes) are part of community 7, while the entire community only represents 4.1% of the network. Community 7 includes 20.5% of putative recognition genes, 37.9% of modulation genes, and 26.7% of effector genes, but only 1 (TOLL5D) of 53 annotated immune signal transduction genes. The largest immune protein family to be overrepresented is the CLIP-domain containing serine proteases (CLIPs) [79], with 37 of 88 of annotated CLIPs are present in Community 7.

In addition to higher-order clustering of biological processes within communities, we also compared enriched GO terms between communities. We found that in many instances, related biological processes are enriched in neighboring communities. For example, Communities 3, 10, and 13 map to the same region of the 2D visualized network (Fig. 12) and are enriched for genes with GO term annotations related to chitin metabolism. These GO terms are partially explained by the enrichment of Communities 3 and 10 for genes encoding CPR proteins, which are characterized by a Rebers and Riddiford Consensus (RR) domain and are major components of the insect cuticle. CPR proteins fall into two evolutionary subgroups, based on their RR domain type, which are referred to as RR1 and RR2. The *An. gambiae* genome encodes 55 RR1 and 102 RR2 genes [80], of which 47 RR1 and 92 RR2 genes are present in the AgGCN1.0. Community 3 contains 45% of CPR genes, while only containing 13.6% of genes in the network. Community 13 contains 19% of RR1 genes, while only containing 1.0% of genes in the network.

Other examples are of shared GO terms across neighboring communities are the enrichment of innate immunity genes in communities 7 and 9, as well as the enrichment of salivary gland protein genes in communities 15 and 9 (Fig. 11). The algorithm of the Force Atlas 2.0 network layout pulls together nodes that are connected by links, while repelling nodes [70], thus the expression of genes within neighboring communities is likely more similar than to communities that are more distant to each other.

## 6. Conclusion

In this paper, we constructed a global gene co-expression network for *An. gambiae* based on the metaanalysis of 30 gene expression studies. The rich information produced by different experiments made it challenging with existing methodologies to analyze the relations of genes based on co-expression, as different methodologies were used to process the data. Current approaches cannot directly construct a GCN from many conditions that are tested with various methodologies. The raw expression values of each condition therefore required normalization before applying any network construction methods. In this work, we adopted the z-score normalization, by which the expression values in different conditions are normalized with zero mean and unity variance. The co-expression of genes was then quantified by the Pearson correlation coefficient, which is computed for each pair of genes based on the normalized expression. However, the number of paired elements between genes was heterogeneously distributed, given that only a subset of genes was tested under each condition, and missing values are ubiquitous in experimental data. Thus, a unique threshold or criterion was not appropriate for selecting edges. Instead, we categorized the PCCs into different intervals according to the number of paired elements, and the fitted sliding threshold was used to select edges for the GCN. In addition, the PCCs were rescaled by the reversed curve of the fitted sliding thresholds to obtain the edge weights.

Analyses of the resulting AgGCN1.0 network showed that it is robust with respect to random removal of up to 15% conditions. In addition, the sensitivity of the AgGCN1.0 structure to the underlying number of conditions mostly affected the loss of those edges derived from few experimental data. Studies of the topological properties of the network showed that AgGCN1.0 is dominated by hub nodes and approximates a scale-free property. Scale-free properties are typical for real-world networks [81, 82], including GCNs and other biological networks [68, 83]. This property likely provides network robustness and protects against random perturbations, e.g., mutations that protect the global architecture of the network. In addition, the global architecture is also protected against targeted perturbations, due to the node properties of the AgGCN1.0 network core, where all centrality measures are maximized rather than a specific one for each core node. It will be interesting to compare whether this feature is common of global GCNs, and whether it is a consequence of specific biological gene properties or functions.

Scale-free and small-world networks are characterized by the existence of communities, which are groups of nodes that are well-connected insides and loosely connected with other communities. This translates to GCNs, in which genes are often grouped into modules that are characterized by a similar function. The 15 communities of the AgGCN1.0 are indeed enriched for different GO and KEGG terms, thus constituting biologically meaningful groups. This finding provides additional validation of the updated GCN construction methodology reported in this study. Not surprisingly, many communities were enriched functions that constitute fundamental biological processes required for cellular function regardless of cell or tissue type. This is largely explained by the non-model organism status of mosquitoes. With limited functional characterization of lineage-specific genes, gene annotation often relies on orthology and thus biasing even further the inherent incompleteness of gene ontology [84]. While mosquito-specific GO terms have been defined based on anatomy [85], several physiologies highly relevant to mosquito biology are either not included in the GO database (e.g., hematopaghy) or are underutilized in annotations (e.g., host-seeking). Nevertheless, community structure in the AgGCN1.0 is clearly defined by functional clustering of genes.

The GO term and KEGG pathway enrichment analysis also revealed an interesting pattern that may extend beyond the AgGCN1.0 to other gene co-expression networks. The architecture of the AgGCN1.0 at its *s*-core is defined by genes required for respiration and translation, both fundamental processes required for survival at the cellular level. The expression of these genes is further integrated into the expression of genes required for aerobic energy production and cellular protein genesis and homeostasis. Not surprisingly, these genes are well-conserved evolutionarily at the metazoa and arthopoda level [86]. This not only demonstrates integration of these processes at the transcriptional level, but also perhaps that the transcriptional regulation of these genes evolved early and has been maintained throughout evolution. In contrast, communities at the periphery of the network (e.g., communities 11 and 15) are enriched in genes that contribute to specialized biological processes, including blood feeding [77] and potentially sensory neuron and/or sperm function [87]. Genes in these communities also tended to be more lineage-specific, suggesting that integration of novel biological processes into the *An. gambiae* transcriptome may not require rewiring of the regulatory circuity. In the future, it will be interesting to determine whether gene age is a principle that governs global gene co-expression patterns in *An. gambiae* and beyond.

In summary, this manuscript provides a correlation-based methodology to build GCNs from highly heterogeneous expression data. This methodology was then applied successfully to build a robust global GCN for *An. gambiae*, updating the previous meta-analysis performed by [12]. This network is available freely to the scientific community at https://ece.k-state.edu/netse/projects/sprojects/NIHproj.html. Analysis of the AgGCN1.0 LCC suggests that the architecture of the *An. gambiae* transcriptome maximizes integration of essential cellular processes and enables evolutionary flexibility to integrate the expression of novel biological functions.

## Acknowledgment

We thank Dr. Robert MacCallum, Imperial College London, UK for initial advice and sharing of the data set Anopheles-gambiae EXPR-STATS VB-2019-02, which used to be publicly available through VectorBase (www.vectorbase.org). This work has been supported by the National Institutes of Health under Grant No. R01AI140760 (KM) and R01AI148529 (NB). This is contribution no. 22-162-J from the Kansas Agricultural Experiment Station. The contents of this article are solely the responsibility of the authors and do not necessarily represent the official views of the funding agencies.

